# Hetero-antagonism of avibactam and sulbactam with cefiderocol in carbapenem-resistant *Acinetobacter* spp

**DOI:** 10.1101/2024.03.04.583376

**Authors:** Olivia Wong, Vyanka Mezcord, Christina Lopez, German Matias Traglia, Fernando Pasteran, Marisel R. Tuttobene, Alejandra Corso, Marcelo E. Tolmasky, Robert A. Bonomo, María Soledad Ramirez

**Affiliations:** Center for Applied Biotechnology Studies, Department of Biological Science, College of Natural Sciences and Mathematics, California State University Fullerton, Fullerton, California, USA; Unidad de Genómica y Bioinformática, Departamento de Ciencias Biológicas, CENUR Litoral Norte, Universidad de la República, Uruguay; Laboratorio Nacional/Regional de Referencia en Antimicrobianos, Instituto Nacional de Enfermedades Infecciosas, ANLIS Dr. Carlos G. Malbrán, Buenos Aires, Argentina; Área Biología Molecular, Facultad de Ciencias Bioquímicas y Farmacéuticas, Universidad Nacional de Rosario, Rosario, Argentina; Instituto de Biología Molecular y Celular de Rosario (IBR, CONICET-UNR), Rosario, Argentina; Research Service and GRECC, Louis Stokes Cleveland Department of Veterans Affairs Medical Center, Cleveland, Ohio, USA; Departments of Medicine, Pharmacology, Molecular Biology and Microbiology, Biochemistry, Proteomics and Bioinformatics, Case Western Reserve University School of Medicine, Cleveland, Ohio, USA; CWRU-Cleveland VAMC Center for Antimicrobial Resistance and Epidemiology (Case VA CARES), Cleveland, Ohio, USA

**Keywords:** *Acinetobacter*, antagonism, synergy, cefiderocol, antimicrobial susceptibility testing (AST), diazabicyclooctanes (DBOs), carbapenem-resistance, NDM

## Abstract

The emergence of Gram-negative bacteria resistant to multiple antibiotics, particularly carbapenem-resistant (CR) *Acinetobacter* strains, poses a significant threat globally. Despite efforts to develop new antimicrobial therapies, limited progress has been made, with only two drugs—cefiderocol and sulbactam-durlobactam—showing promise for CR-*Acinetobacter* infections. Cefiderocol, a siderophore cephalosporin, demonstrates promising efficacy in the treatment of Gram-negative infections. However, resistance to cefiderocol has been reported in *A. baumannii*. Combination therapies, such as cefiderocol with avibactam or sulbactam, show reduced MICs against cefiderocol-non-susceptible strains with in vivo efficacy, although the outcomes can be complex and species-specific. In the present work, the molecular characterization of spontaneous cefiderocol-resistant variants, a CRAB strain displaying antagonism with sulbactam and an *A. lwoffii* strain showing antagonism with avibactam, were studied. The results reveal intriguing insights into the underlying mechanisms, including mutations affecting efflux pumps, transcriptional regulators, and iron homeostasis genes. Moreover, gene expression analysis reveals significant alterations in outer membrane proteins, iron homeostasis, and β-lactamases, suggesting adaptive responses to selective pressure. Understanding these mechanisms is crucial for optimizing treatment strategies and preventing adverse clinical outcomes. This study highlights the importance of preemptively assessing drug synergies to navigate the challenges posed by antimicrobial resistance in CR-*Acinetobacter* infections.

## Introduction

The widespread dissemination of Gram-negative bacteria displaying resistance to virtually all available antibiotics raises significant concerns (1, 2). Within this context, the WHO and the Centers for Disease Control and Prevention (CDC) have recently categorized carbapenem-resistant (CR) *Acinetobacter* as a critical priority pathogen (2, 3). The global emergence and widespread prevalence of *Acinetobacter* strains resistant to multiple antibiotic classes underscore the pressing demand for new antimicrobial therapies (1, 3, 4). Despite significant efforts by diverse research entities and pharmaceutical companies over the past decade (5–7), the approval of novel drugs specifically targeting CR-*Acinetobacter* has been limited. Presently, only two drugs—cefiderocol and sulbactam-durlobactam—have received approval from the U.S. Food and Drug Administration (FDA) for treating *Acinetobacter baumannii-calcoaceticus* complex infections. (https://www.accessdata.fda.gov/drugsatfda_docs/label/2019/209445s000lbl.pdf, https://www.accessdata.fda.gov/drugsatfda_docs/label/2023/216974Orig1s000Correctedlbl.pdf).

Cefiderocol, a siderophore cephalosporin, possessed a chlorocatechol group situated at the termination of the C3 side chain that chelates iron (Fe^3+^) and enhances its stability against β-lactamase (8, 9). This feature enables the molecule to efficiently pass through the outer cell membrane of Gram-negative bacteria (GNB) into the periplasmic space of GNB using the Ton-B dependent iron transport system (10).

The Infectious Diseases Society of America recommends combination therapy for treating CR-*Acinetobacter baumannii* (CRAB) infections, especially regimens incorporating sulbactam (11). However, the clinical evidence supporting this remains controversial (12). Cefiderocol should be reserved for cases where other antibiotics fail, or resistance limits their use. When prescribing cefiderocol for CRAB infections, it’s suggested as part of a combination therapy regimen (11).

Recent treatment developments include combining either sulbactam or avibactam with cefiderocol, resulting in reduced minimum inhibitory concentrations (MICs) against cefiderocol-non-susceptible (13). In vivo models demonstrate efficacy for these combinations against all evaluated cefiderocol-non-susceptible CRAB isolates (14). Avibactam, a non-β-lactam β-lactamase inhibitor, addresses resistance to cefiderocol by suppressing cefiderocol-hydrolyzing secondary enzymes like VEB and PER, potentially present in CRAB isolates (15). Sulbactam, known for weakly inhibiting intrinsic ADCs, could provide activity in this combination via ADC inhibition with its high exposure (14).

Our prior research shows various DBO inhibitors enhance cefiderocol’s in vitro effectiveness against CRAB, with notable effects at higher cefiderocol MICs (13). Unlike other studies, combining cefiderocol with sulbactam showed inconsistent results, with only a 20% enhancement noted. In this study, we delve into the molecular characterization of two CR-*Acinetobacter* isolates that exhibited unanticipated antagonistic behavior against cefiderocol: a CRAB strain displaying antagonism with sulbactam, and an *A. lwoffii* strain showing antagonism with the combination of cefiderocol plus avibactam. The insights gained from this research help unravel the rationale behind the unexpected antagonistic phenomenon and it serves as a vital warning about the importance of pre-evaluating these synergies to prevent a potential adverse impact on clinical outcomes.

## Material and Methods

### Bacterial strains

The carbapenem-resistant clinical *Acinetobacter* isolates AMA22_1 (*A. baumannii)* and AMA23 (*A. lwoffii*), both containing *bla*_NDM-1_, were used in this study (13, 16). In addition, spontaneous cefiderocol resistant variants (AMA22_1 IHCA and AMA23 IHC AVI 4 μg/mL and R AVI 8 μg/mL of the AMA22_1 and AMA23 parental strains, which have emerged within the inhibition ellipse zones of cefiderocol in cation-adjusted Mueller-Hinton agar (CAMHA) containing 4 ug/ml of sulbactam or 4 to 8 ug/ml avibactam, respectively, were also use (Fig. 1).

**Fig. 1.**
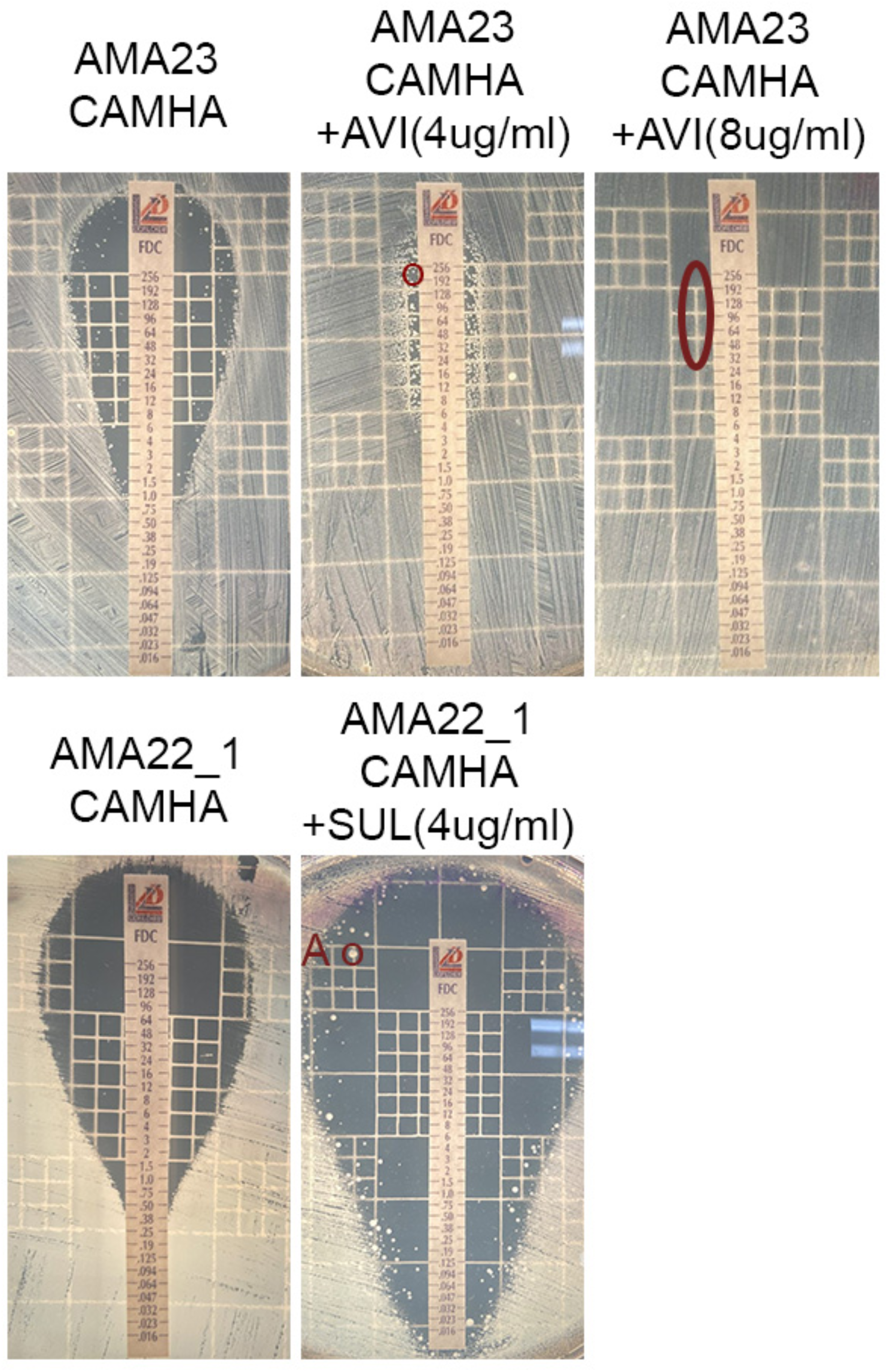
Effect of β-lactamase inhibitors (avibactam (AVI) or sulbactam (SUL)) on the antimicrobial susceptibility of *Acinetobacter* spp. to performed cefiderocol susceptibility. Minimum inhibitory concentration (MIC) was performed following manufacturer’s recommendations (Liofilchem S.r.l., Italy). Red circles indicate intracolonies chosen for this study (AMA22_1 IHC A, AMA23 IHC AVI 4 μg/mL, and AMA23 R AVI 8 μg/mL).

The naturally mutant variants of both strains showed different levels of cefiderocol susceptibility. The selected mutant variants were stored at -80°C as Luria Bertani (LB) broth containing 20% glycerol stocks. The stability of the different levels of cefiderocol resistance in the mutant variants was performed by daily subcultures in antibiotic free plates.

### Antimicrobial susceptibility testing (AST)

The antibiotic susceptibility assays were carried out in accordance with the Clinical and Laboratory Standards Institute (CLSI) guidelines (17). AMA22_1, AMA23 and its corresponding variants cells were cultivated in CAMHA or iron-depleted CAMHB, when corresponded, and adjusted to a 0.5 McFarland standard value, were utilized for the assays. Disks and commercial E-strips (Liofilchem S.r.l., Roseto degli Abruzzi, Italy) containing varying concentrations of different antibiotics and were employed. These included imipenem (IMI), meropenem (MRP), ceftazidime (CAZ), ceftriaxone (CRO), gentamicin (GM), ciprofloxacin (CIP), ceftazidime-avibactam (CZA), cefiderocol (FDC), ampicillin (AMP), ampicillin-sulbactam (AMS), and amikacin (AN). Cefiderocol MICs were also performed by two different dilution methods, comASP (Liofilchem S.r.l., Roseto degli Abruzzi, Italy) and in house broth microdilution (BMD) with ID-CAMHB. All commercial assays strictly followed the manufacturer’s instructions (https://www.liofilchem.com/images/brochure/mic_test_strip_patent/MTS51.pdf).

Additionally, in specific cases, CAMHA medium was supplemented with 1, 2, 4, 8 and 16 μg/mL of sulbactam or avibactam (Sigma-Aldrich). Measurements were taken after incubating the plates at 37°C for 18 hours. Interpretation of the data was done based on CLSI breakpoints (17), with *Escherichia coli* ATCC 25922 employed for quality control purposes. Each susceptibility assay was repeated at least twice using independent biological samples on each occasion.

### Whole genome sequence analysis

The genomic DNA extraction of the parental strains (AMA22_1 and AMA23) and the three randomly selected variants (AMA22_1 IHCA and AMA23 IHC AVI 4 μg/mL and R AVI 8 μg/mL) was performed following the manufacturer’s instructions using the Wizard Promega kit (Promega, Madison, WI, USA). For whole genome sequencing, SEQCENTER sequencing service (Pittsburgh, PA, USA) performed the outsourced procedure utilizing NextSeq 550 Illumina technology. To ensure sequence quality, FASTQC software analysis (https://www.bioinformatics.babraham.ac.uk/projects/fastqc/) was conducted, followed by trimming and filtering using Trimmomatic software (version: 0.40, ILLUM-NACLIP: TrueSeq3-PE.fa.2:30:10; LEADING:3; TRAILING:3; SLIDINGWINDOW: 4:15; MINLEN:36) (18). De novo sequence assembly was carried out using SPAdes (version: 3.15.4, default parameters) (19) and subsequently assessed for quality using QUAST (version: 5.2.0) (20). Genome annotation was accomplished via PROKKA (21), whereas variant calling employed the *breseq* and *gdtools* software packages (version: 0.38.1, consensus mode, default parameters) (22). Recombination regions were detected and eliminated using Gubbins software (version: 3.3.0, default parameters) (23). The raw genomic sequencing data and assemblies have been deposited in the Zenodo repository (https://zenodo.org/records/10729558).

### RNA extraction and transcriptional analysis through RT-qPCR

For RNA extractions, overnight cultures of AMA22_1, AMA23, AMA22_1 IHCA, AMA23 IHC AVI 4 μg/mL, and R AVI 8 μg/mL strains underwent a 1:10 dilution in iron-depleted CAMHB and were incubated with agitation at 200 rpm for 18 hours at 37°C. RNA extraction from each sample utilized the Direct-zol RNA Kit (Zymo Research, Irvine, CA, USA) following the manufacturer’s protocol. RNA extractions were done in triplicates. Subsequently, the DNase-treated RNA was used to synthesize cDNA employing the iScriptTM Reverse Transcription Supermix for qPCR reagents (Bio-Rad, Hercules, CA, USA) following the provided manufacturer’s guidelines. The concentration of the resulting cDNA was adjusted to 50 ng/μL and 2 μL of the adjusted cDNA was used to carry out qPCR reaction using qPCRBIO SyGreen Blue Mix Lo-ROX according to the manufacturer’s instructions (PCR Bio-systems, Wayne, PA, USA).

In Table S1, the list of the primers used for the transcriptional analysis are listed. Each cDNA was tested independently in triplicate utilizing the CFX96 TouchTM Real-Time PCR Detection System (Bio-Rad, Hercules, CA, USA). The transcript levels of each sample were standardized against the *rpoB* transcript levels in individual cDNA samples (24). The quantification of gene expression was conducted using the comparative threshold method 2^-ΔΔCt^ (25). Statistical differences were determined using ANOVA followed by Tukey’s multiple comparison test (*P* < 0.05) employing GraphPad Prism (GraphPad software, San Diego, CA, USA).

## Results and Discussion

### Spontaneous occurrence of cefiderocol resistance in the presence of β-lactamase inhibitors

When testing the combination of cefiderocol with β-lactamase inhibitors, such as avibactam, relebactam, zidebactam, and sulbactam, the occurrence of colonies within the halo of inhibition were observed for two CR_*Acinetobacter* strains (AMA22_1 and AMA23) when sulbactam or avibactam were present at a concentration of 4 μg/ml in the CAMHA, respectively (13). For the present study we randomly selected spontaneous resistant variants of AMA22_1 and AMA23 (AMA22_1 IHCA and AMA23 IHC AVI 4 μg/mL) for further studies (Fig. 1A). In addition, for the AMA23 strain a confluent cefiderocol resistance (>256 μg/mL) was noted when testing increases concentrations of avibactam. Cells from this plate were selected for WGS (AMA23 R AVI 8 μg/mL) (Fig. S1).

Cefiderocol MIC determinations were conducted using three different methods (E-strips, comASP, and broth microdilution assays), revealing that all three selected strains exhibited higher levels of resistance compared to their corresponding parental strains (Table 1). In addition, to check if the occurrence of spontaneous cefiderocol resistant colonies was dependent of the β-lactamase inhibitor concentration, increasing concentrations of sulbactam or avibactam were tested for AMA22_1 and AMA23, respectively. As expected, and confirming previous observations, colonies within the zone of inhibition were observed within the inhibition ellipse with 4 μg/mL of sulbactam and avibactam for AMA22_1 and AMA23, respectively (Fig. S1). In addition, for AMA23, as mentioned above, MIC values of > 256 μg/mL were seen when 8 μg/mL of avibactam used to supplement the CAMHA (Fig. S1 and Table 2). In vitro assessment revealed a paradoxical effect of avibactam concentration on cefiderocol in AMA-23, demonstrating synergy at concentrations below 2 mg/L or above 16 mg/L, while exhibiting antagonism at concentrations near the breakpoint. Evaluation of increasing sulbactam concentrations was limited to values <= 8 mg/L due to observed difficulty in AMA22_1 growth beyond this threshold, implying values nearing or surpassing the MIC. Nonetheless, at lower concentrations, a similar paradoxical effect was observed, displaying antagonism near the breakpoint. This indicates a concentration range that could prompt an *Acinetobacter* adaptation to combinations of cefiderocol with inhibitors.

**Table 1.**
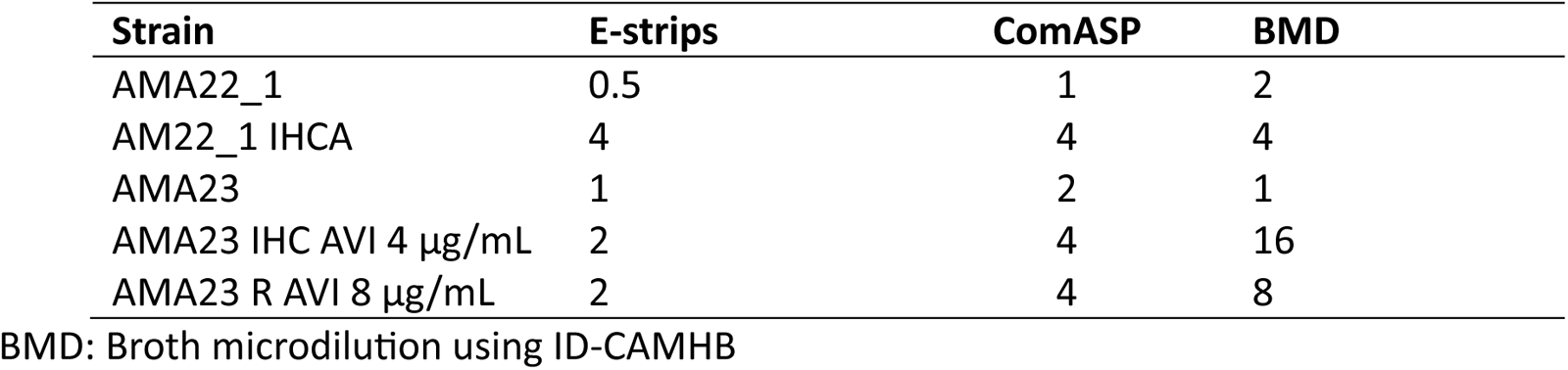
Cefiderocol Minimum inhibitory concentration (MIC) of *A. baumannii* AMA22_1, *A. lwoffii* AMA23 and the isogenic mutant variants obtained within the inhibition ellipse zones of cefiderocol containing 4 ug/ml of sulbactam or 4 to 8 ug/ml avibactam, respectively using gradient strips and microdilution assays.

| Strain | E-strips | ComASP | BMD |
| --- | --- | --- | --- |
| AMA22_1 | 0.5 | 1 | 2 |
| AM22_1 IHCA | 4 | 4 | 4 |
| AMA23 | 1 | 2 | 1 |
| AMA23 IHC AVI 4 µg/mL | 2 | 4 | 16 |
| AMA23 R AVI 8 µg/mL | 2 | 4 | 8 |

BMD: Broth microdilution using ID-CAMHB

**Table 2.** Cefiderocol Minimum inhibitory concentration (MIC) of AMA22_1 and AMA23 in cation-adjusted Mueller Hinton Agar (CAMHA) containing increases concentration of sulbactam or avibactam, respectively using gradient strips.

| Cefiderocol MICs (E-strips) |  |  |  |
| --- | --- | --- | --- |
| Drug and cc (ug/ml) | AMA22_1 | Drug and cc (ug/ml) | AMA23 |
| CAMHA 0 µg/mL SUL | 0.5 | CAMHA 0 µg/mL AVI | 1* |
| CAMHA + 1 µg/mL SUL | 0.19 | CAMHA + 1 µg/mL AVI | 0.5 |
| CAMHA + 2 µg/mL SUL | 0.047 | CAMHA + 2 µg/mL AVI | 0.75 |
| CAMHA + 4 µg/mL SUL | 48* | CAMHA + 4 µg/mL AVI | >256 |
| CAMHA + 8 µg/mL SUL | 0.023 | CAMHA + 8 µg/mL AVI | >256 |
| CAMHA + 16 µg/mL SUL | <0.016 | CAMHA + 16 µg/mL AVI | 0.5 |

SUL= sulbactam, AVI= avibactam, \*occurrence of intracolony within the inhibition ellipse zones of cefiderocol.

Additionally, there is no loss of the enhanced resistance phenotype after 10 daily subcultures, which indicates that the resistance phenotype is a stable trait.

The susceptibility of AMA22_1 IHCA, AMA23 IHC AVI 4 μg/mL, and AMA23 R AVI 8 μg/mL to various antibiotics (including meropenem, imipenem, gentamicin, ampicillin/sulbactam, amikacin, ciprofloxacin, levofloxacin, tigecycline, colistin, and trimethoprim-sulfamethoxazole) was evaluated to identify any possible collateral-sensitivity or cross-resistance to cefiderocol. The susceptibility results of all the strains are showed in Table S2. No differences in resistance profiles were observed between the parental strain and the corresponding variants for most of the antibiotics tested. Only, collateral susceptibility to ampicillin and amikacin were observed for AMA23 variants (Table S2), which was confirmed by MIC determination showing a 2-fold and 4-fold decrease dilution, respectively (data not shown). No differences in resistant profiles were observed between the parental strain and the corresponding variants.

Between 20-80% of *Acinetobacter* clinical isolates displayed heteroresistance to cefiderocol, defined as the presence of heterogeneous bacterial populations wherein subpopulations exhibit higher levels of antibiotic resistance than the main population (26). Exploration of heteroresistance to ampicillin-sulbactam has been limited to one study, revealing its presence in up to 25% of isolates in ICU patients (27).

While the clinical significance of this phenomenon remains debated, recent research has shed light on key aspects of cefiderocol-heteroresistance. Notably, observations of small colonies within inhibition zones reverting to their original form upon drug removal suggest the instability of cefiderocol-heteroresistance. Additionally, strains harboring multiple classes of β-lactamase resistance genes may see restored activity with the addition of β-lactamase inhibitors like ceftazidime/avibactam to cefiderocol (28). Moreover, animal models have shown that combining ceftazidime-avibactam can impede the development of in vivo resistance to cefiderocol, while in vitro, microcolonies within the inhibition zone notably decreased for *A. baumannii* isolates with high-end susceptibility (MIC 2 mg/L) (14).

In our study, we observed that heteroresistant colonies maintained their resistant trait after serial passage in antibiotic-free media, and inhibitors at typical assay concentrations did not reverse the resistance. Most of these studies have primarily examined *A. baumannii*. Expanding our investigation to encompass a group of CR-*Acinetobacter* species distinct from *A. baumannii*, we observed a contrasting phenomenon: while most isolates displayed no alteration or an elevation in MIC values of cefiderocol when tested with β-lactamase inhibitors, a notable synergy was evident with sulbactam (13). These findings suggest species-specific variations in response to combination therapies.

### Distinct changes at the genome level of the selected spontaneous resistant variants

The analysis of the nucleotide substitutions of AMA22_1 IHCA revealed eight synonymous and three non-synonymous mutations. Of the non-synonymous mutations, two affected genes encoding hypothetical proteins, while one impacted the gene encoding the transposase IS*Aba26* (Table S3). Additionally, a deletion of two nucleotides was identified in the gene encoding a hypothetical protein, and an insertion of two nucleotides was found in the gene encoding the Cobalt-zinc-cadmium resistance protein CzcA. These genes are components of an efflux pump belonging to the RND group, a superfamily that includes various efflux pumps associated with multidrug resistance phenotypes, such as the AdeABC efflux pump. While CzcA is a part of the CzcCBA efflux pump (29), which confers resistance to heavy metals, its role in multidrug resistance remains unexplored. Investigating the association of CzcCBA with multidrug resistance will be crucial for understanding its function in this context. None of intergenic changes were found in regulator or promoter regions (Table S3).

The analysis of the nucleotide substitutions of AMA23 IHC AVI 4 μg/mL, and AMA23 R AVI 8 μg/mL revealed 18 and15 synonymous mutations, respectively.

Three non-synonymous mutations were detected, and notably, these mutations were identical in both isolates. They affected genes responsible for encoding the BfmR DNA-binding transcriptional regulator and two transposases (IS*66* and IS*Aba31*) (Table S3). BfmR has been recognized for its role in regulating biofilm formation and has also been associated with enhancing meropenem resistance (30–32). These mutations not only have the potential to influence meropenem (33) resistance but may also impact a wide range of beta-lactam antibiotics, including cefiderocol.

An insertion of 17 nucleotides was identified in a gene encoding a transposase (IS*Aba125*). Mutations within insertion sequences can affect the functionality of downstream genes’ promoters/enhancers and the transposition mechanism (34). However, none of the insertion sequences harboring mutations were observed to affect genes related to iron uptake/homeostasis or resistance to β-lactams and cefiderocol. Additionally, 19 and 23 intergenic changes were found in AMA23 IHC AVI 4 μg/mL and AMA23 R AVI 8 μg/mL, respectively. None of intergenic changes were found in regulator or promoter regions (Table S3).

### Modifications in the transcript levels of key genes in the spontaneous resistant variants

mRNA extraction was performed on AMA22_1 and AMA23, as well as IHC cells, to assess the expression levels of genes related to outer membrane proteins, TonB-dependent transporters and iron homeostasis, the two-component system BaeRS, β-lactamase, and *pbp3*. In Figure 2A, the results indicate significant down-regulation of *ompA*, *carO*, β-lactamase genes, and *pbp3* in the AMA22_1 IHCA compared to the parental strain. Conversely, genes encoding for BaeRS exhibited significant up-regulation in AMA22_1 IHCA. Interestingly, out of the four iron-related genes studied, *pirA* and *ppiA* were found to be downregulated, while increased expression levels were observed fo*r bauA* and *piuA* (Fig. 2A and Table S4). In AMA23 cells, the genes encoding outer membrane proteins (*ompA* and *carO*) showed down-regulation in both mutant variant strains (Fig. 2B and Table S4). Notably, an increased expression of *bla*_NDM-1_ was observed in AMA23 IHC AVI (Fig. 2B and Table S4). There were no significant differences noted in the TonB-dependent transporters and iron homeostasis for AMA23 cells, except for *piuA*, which exhibited significant up-regulation in AMA23 R AVI 8 μg/mL (Fig. 2B and Table S4). A significant enhancement of the transcript levels of the gene encoding for the Two-component system BaeSR was seen in AMA23 R AVI 8 μg/ml (Fig. 2B and Table S4).

**Fig. 2.**
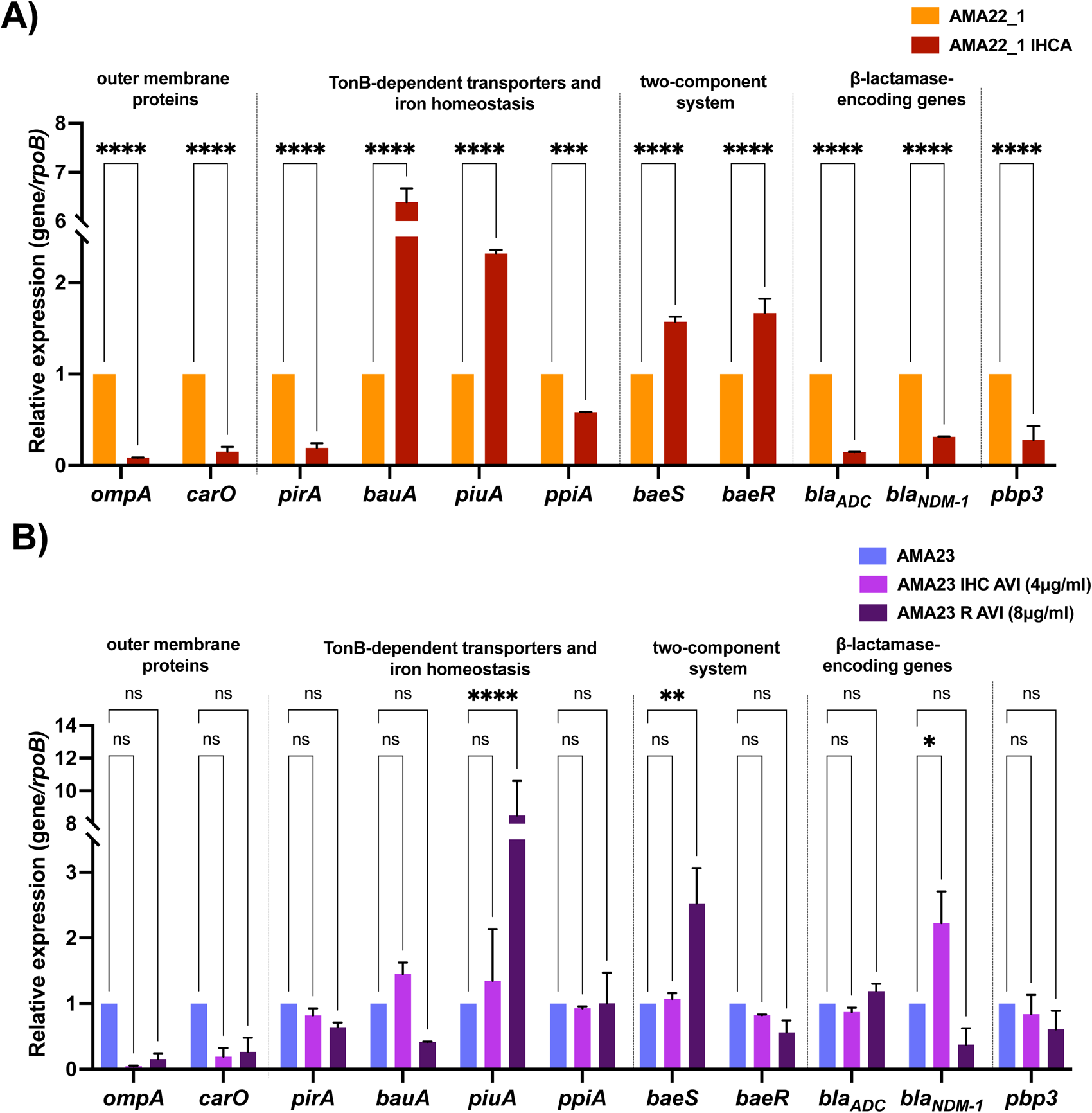
Expression of genes coding for outer membrane proteins (*ompA* and *carO*), TonB-dependent transporters and iron homeostasis (*pirA, bauA, piuA* and *ppiA*), the BaeRS two-component system, β-lactamase and *pbp3* in the AMA22_1 and AMA22_1 IHC A (A) or AMA23, AMA23 IHC AVI 4 μg/mL, and AMA23 R AVI 8 μg/mL (B) strains. The data shown of qRT-PCR are mean ± SD. Fold changes were calculated using double ΔCt analysis. At least three independent biological samples were tested using four technical replicates. Statistical significance (p < 0.05) was determined by two-way ANOVA followed by Tukey’s multiple comparison test. Significance was indicated by: * *p* <0.05, ** *p* < 0.01, *** *p* < 0.001 and **** *p* < 0.0001.

As one of the main mutations in the AMA23 variant cells affected the gene that encoding the transcriptional regulator BfmR, the transcriptional expression levels of BfmRS encoding genes in the AMA23 IHC AVI (4μg/ml), AMA23 R AVI (8μg/ml) and its parental strain were evaluated (Fig. S2 and Table S4). Differences were no observed in the level expression of *bfmS.* On the other hand, *bfmR* were significantly enhance in both AMA23 IHC AVI 4 μg/mL, and AMA23 R AVI 8 μg/mL (Fig. S2 and Table S4).

Previous reports have indicated significant alterations in the expression levels of genes related to iron-uptake systems, β-lactam resistance, and biofilm formation in heteroresistant variants of a CRAB strain, particularly in cultures supplemented with cefiderocol and human fluids (32).

Additionally, mutations in the *baeSR* genes were associated with reduced susceptibility of *A. baumannii* to cefiderocol by up-regulating the expression of the MFS family efflux pump and MacAB-TolC efflux pump (35). The modified expression of these genes seen in the studied heteroresistant cells, AMA22_1 IHCA, AMA23 IHC AVI, and AMA23 R AVI 8 μg/mL, could potentially be linked to the increased cefiderocol MIC seen in these strains.

The complexity of the regulation of gene expression known to contribute to cefiderocol resistance in different *Acinetobacter* species and mutant variants can explain the different levels observed for some of the genes evaluated. However, several genes implicated in bacterial permeability, such as *ompA* and *carO*, along with genes associated with TonB iron uptake systems, such as *pirA*, and two-component system consistently exhibited a similar pattern across all studied variants, contributing to resistance to cefiderocol.

### Concluding remarks

This study underscores the complex nature of emergent resistance to cefiderocol in *Acinetobacter* spp. Unlike the acquired resistance to some antibiotics, which is typically mediated by the horizontal acquisition of one or more genes, the resistance in the strains described in this work is associated with numerous genomic mutations and phenotypic changes in the expression of various genes. These findings suggest that resistance may result from multiple and heterogenous mechanisms. Moreover, the antagonistic effect observed when cefiderocol is used in combination with avibactam or sulbactam suggests a myriad of intricate interactions resulting from the mentioned genotypic and phenotypic modifications. Such interactions indicate a complex network of interactions that can affect the efficacy of combination therapies containing cefiderocol, thereby complicating the prediction of therapeutic outcomes. Therefore, a prior evaluation of synergies or antagonisms using specific antibiotics on a case-by-case basis is essential to prevent potential undesired clinical outcomes.

## Author Contributions

F.P., O.W. V.M., C.L., G.M.T., and M.S.R. conceived the study and designed the experiments. F.P., M.R.T., A.C., M.E.T., R.A.B, and M.S.R. analyzed the data and interpreted the results. F.P., M.R.T., A.C., M.E.T., and M.S.R. contributed reagents/materials/analysis tools. F.P., G.M.T., A.C., M.E.T., R.A.B, and M.S.R wrote and revised the manuscript. All authors read and approved the final manuscript.

